# Wnt5A supports antigen cross-presentation and CD8 T cell activation

**DOI:** 10.1101/2022.01.04.474906

**Authors:** Tresa Rani Sarraf, Malini Sen

## Abstract

Antigen processing, cross-presentation, and antigen-specific CD8 T cell response form part and parcel of T cell-mediated immunity. Yet, lacunae remain in our understanding of antigen processing/presentation and CD8 T cell response. Given the association of Wnt5A signaling with immune homeostasis, we evaluated the utility of Wnt5A in antigen processing, cross-presentation, and CD8 T cell activation. Using mouse bone marrow-derived dendritic cells as antigen-presenting cells and ovalbumin as a model antigen we found that Wnt5A mediated regulation of actin and proteasome dynamics is inherently associated with antigen processing. A Wnt5A-Actin-Protasome axis also contributes to antigen cross-presentation and antigen responsive CD8 T cell expansion. In concurrence with these observations, we demonstrated impaired activation of ovalbumin-specific CD8 T cells in ovalbumin immunized Wnt5A heterozygous mice as illustrated by their poor CD8 T cell recall response to ovalbumin when compared to similarly immunized wild type cohorts. Our results suggest that Wnt5A signaling-directed antigen processing/presentation could be vital for generating CD8 T cell recall response to antigen, thus shedding light on a critical parameter of immunity.

## Introduction

Antigen processing, its cross-presentation to CD8 T cells, and antigen-specific CD8 T cell expansion constitute a vital feature of cell-mediated immunity to viral and other microbial infections. Quite naturally, impairments in any of these processes could lead to increased susceptibility to infection. However, several lacunae remain in our understanding of how host intracellular processes are coordinated toward the activation and preservation of antigen-specific CD8 T cell responses. (Embgenbroich and Burgdorf, 2018; Huang et al., 2014; Velilla et al., 2006)

Contemporary literature suggests that Wnt5A signaling is important for the maintenance of cell polarity and organelle dynamics (Gon et al., 2013; Witze et al., 2008). Thus it is not surprising that Wnt5A signaling is also a major regulator of the cytoskeletal actin network, which constitutes the fundamental framework for organelle crosstalk (Kjeken et al., 2004; Oberhofer et al., 2020; Semenova et al., 2008). We previously demonstrated that Wnt5A signaling is required for pathogen killing through phagocytosis and xenophagy, which are inherently linked with modulations in actin assembly (Chakraborty et al., 2017; Jati et al., 2021, 2018). Antigen processing and its presentation toward T cell activation being a functional outcome of actin remodeling and associated intracellular organelle trafficking (Liu et al., 2013), it was important to investigate if Wnt5A signaling promotes antigen processing and presentation, thereby controlling T cell activation.

Wnt5A belongs to a large family of secreted Wnt glycoprotein ligands that signal upon binding to Frizzled and/or ROR cell surface receptors (Cadigan and Nusse, 1997; Nusse, 2005). Wnt-Frizzled/ROR signal transduction occurs in various cell types ranging from epithelial and mesenchymal cells to specialized cells of the immune system including lymphocytes, macrophages and dendritic cells (Gon et al., 2013; Malsin et al., 2019; Okoye et al., 2008; Valencia et al., 2014; van Loosdregt et al., 2013). On account of sequence homology among the different Wnt ligands and their receptors, these signaling entities function in multiple combinations under different physiological conditions. The mode of Wnt-Frizzled/ROR signaling in a particular cell type in fact depends to a great extent on the existing ligand–receptor stoichiometry and the availability of signaling intermediates (Mikels and Nusse, 2006). While the canonical mode of Wnt signaling is commonly associated with the transcriptional coactivator β-catenin, the non-canonical mode, of which Wnt5A signaling is a representative, is able to operate independent of β-catenin; although crosstalk with canonical signaling intermediates is not uncommon (Sheng et al., 2014).

In the current study, using ovalbumin (OVA) as a model foreign antigen and mouse bone marrow-derived dendritic cells (BMDC) as antigen-presenting cells we demonstrated that Wnt5A signaling promotes antigen processing and presentation to CD8 T cells leading to CD8 T cell activation. Furthermore, we depicted that the occurrence of partial Wnt5A gene silencing in Wnt5A heterozygous (Wnt5A+/-) mice correlates with considerably subdued OVA-specific CD8 T cell recall response, suggesting that Wnt5A signaling dependent antigen processing/presentation and CD8 T cell activation may be crucial for the sustenance of antigen-specific CD8 T cell memory.

## Materials and Methods

Reagents: RPMI 1640 (Cat No.-31800-022), Fetal Bovine Serum (FBS: Cat no.-10082147), PenStrep (Cat no.-15140122), and L-Glutamine (Cat no.-25030081) for cell culture were purchased from Invitrogen. GM-CSF (Cat no.-315-03) and IL4 (Cat no.-214-14) were purchased from Peprotec. APC-hamster anti-mouse CD11c (Cat No.-550261), Propidium iodide (PI) (Cat No.-51-66211E), PE-mouse anti-mouse H-2K[d] (Cat No.-553566), PE-IgG2a k isotype (Cat No.-553457), Per CP-Cy 5.5-rat anti-mouse I-A/I-E (Cat No.-562363), Per CP-Cy 5.5 rat IgG2b k isotype (Cat No.-550764), V500-Syrian hamster anti-mouse CD3 (Cat No.-560771), BV605-rat anti-mouse CD8 (Cat No.-563152), APC rat anti-mouse CD44 (Cat No.-559250), rat anti-mouse CD62L (Cat No.-553148), APC-R700 rat anti-mouse IL-2 (Cat No.-565186) and PE-CF594-rat anti-mouse IFN-γ were purchased from BD Biosciences. PE-Cyanine 7-rat anti-mouse Granzyme B (Cat. No.-25-8898-82), donkey anti-goat alexa fluor 488 (Cat no.-A-11055), donkey anti-rabbit alexa fluor 546 (Cat no.-A10040) and donkey anti-goat alexa fluor 546 (Cat no.-A-11056) were purchased from eBioscience. Anti-human/mouse Wnt5A monoclonal antibody (Cat no.-MAB645) and anti-rat IgG-HRP (Cat no.-HAF005) were purchased from R & D Systems. Anti-PA28β (Cat no.-BB-AB0190) was purchased from Biobharati Life Sciences and anti-PA28*α* (Cat no.-sc-21267) was purchased from Santa Cruz. DQ-OVA (Cat no.-D12053), FITC-OVA (Cat no.-023020), Prolong glass antifade (Cat no.-P36982), TRIzol reagent (Cat no.-15596018), DAPI (Cat no.-D1306), Alexa fluor 555 phalloidin (Cat no.-A34055), CFSE (Cat no.-C34570), Lipofectamine RNAimax (Cat no.-56532), OptiMEM (Cat no.-31985-070) were purchased from Invitrogen. Poly-l-lysine (Cat no.-P-8920) and Albumin from chicken egg (Cat no.-A5503-1G) were purchased from Sigma. Tween-20 detergent (Cat no.-655205-250ML) and TMB solution (Cat no.-CL07-1000MLCN) were purchased from Merck. Bradford reagent (Cat no.-5000006) was purchased from Biorad. Recombinant Wnt5A (Cat no.-GF146), Western HRP substrate (Cat no.-WBLUC0500), MG132 (Cat no.-474790) were purchased from Millipore. 7-AAD cell viability stain (Cat no.-420403) was purchased from Biolegend. OVA SIINFEKL peptide (257-264) (Cat no.-AS-60193-1) was purchased from Anaspec. Mouse CD8a+ isolation kit (Cat no.-130-104-075) and Mini Separator Columns (Cat no.-130-042-201) were purchased from Miltenyi Biotec. H-2Kb OVA Tetramer-SIINFEKL-PE (Cat no.-TS-5001-1C) was purchased from Medical & Biological Laboratories. Brefeldin (Cat no.-203729-1MG) was purchased from Calbiochem. Purified chicken ovalbumin (OVA, grade VI Cat no: A5503-1G) was purchased from Sigma. Complete (CFA Cat no.-263810) and Incomplete Freund’s Adjuvant(IFA, Cat no.-263910) were purchased from Difco. Wnt5a siRNA (5nmol) (Cat no.-M-065584-00-0005) and control siRNA (5nmol) (Cat no.-D-001206-13-05) were purchased from Dharmacon or Wnt5a siRNA (Cat no.-SR-NP001-001) and control siRNA (Cat no.-SR-CL000-005) were purchased from Eurogentec. cDNA synthesis kit (Cat no.-BB-E0045) and PCR (Cat no.-BB-E0010S) kit were purchased from Biobharati Life Sciences.

### Animal Maintenance

Breeding and maintenance of B6;129S7-*Wnt5a*_*tm1Amc*_/Jmice (Wnt5A+/+ and Wnt5A+/-), purchased from Jackson Laboratory, USA, and BALB/c mice were carried out in institute animal facility. Separation of Wnt5A wild type from Wnt5A heterozygous mice for the experimental purpose was accomplished by genotyping, following Jackson laboratory protocol [https://www.jax.org/Protocol?stockNumber=004758&protocolID=23556]. All animals were maintained in optimum physiological conditions with balanced light and dark cycles in IVC (Individually Ventilated Caging) system with *ad libitum* water and food. All experimental mice were 8-10 weeks old.

### Mice immunization

Mice were immunized following published protocols with some modifications (Aubin et al., 2010). In brief, OVA (100ug) dissolved in 100ul PBS, and an equal volume of CFA or IFA were mixed to form an emulsion and each mouse was injected subcutaneously with the emulsion at the tail base at day 0 (OVA + CFA) and day 14 (OVA + IFA). For the immunization of the B6;129S7-*Wnt5a*_*tm1Amc*_/J mice 20ug SIINFEKL peptide was used with 80ug OVA on account of the complementary MHCI H2-Kb.

### BMDC generation

BMDC were derived from the bone marrow of the femur and tibia of BALB/c mice following published protocols (P. Matheu et al., 2008). In brief, cells collected from the bone marrow were suspended in RPMI supplemented with 10% FBS, 2 mM glutamine,1 μg ml^-1^ streptomycin,1-unit ml^-1^ penicillin, 20 ng/ml mouse recombinant GM-CSF and 10 ng/ml mouse recombinant IL-4, following RBC lysis and washing, and cultured in bacterial plates under normal tissue culture ambience. On the 3^rd^ day, 50% of the culture medium was replaced by a fresh medium and on the 6^th^ day, cells were harvested for experiments.

### BMDC transfection

Wnt5A siRNA and control/scramble siRNA were used for transfecting BMDC using reverse transfection method (Hattori et al., 2017). In brief, for transfection in each well of a 6 well tissue culture plate, siRNA (25nM) and lipofectamine RNAimax were diluted separately in 150μl OptiMEM each and mixed gently. The 300ul mix was incubated for 15 min at room temperature, following which harvested BMDC in OptiMEM (1×10^5^ cells/300ul) were mixed separately with either Wnt5a siRNA-RNAimax or control siRNA-RNAimax in a total volume of 1ml. 1ml cells were plated in each well of a 6-well tissue culture plate. Transfected cell medium was replaced by RPMI supplemented with 10% FBS, 24 hrs post-transfection, and incubation was continued for 24 hr-26 hr until assay. Volume adjustment during transfection was done based on the number of cells transfected.

### RNA isolation and cDNA preparation

RNA isolation from cells was done using the Trizol reagent. cDNA was prepared by the cDNA synthesis kit from BioBharati Life Sciences Kolkata, following instructions provided by the manufacturer. PCR was done using the following pairs of primers: Wnt5a (forward 5’-CAGGTCAACAGCCGCTTCAAC-3’ and reverse 5’-ACAATCTCCGTGCACTTCTTGC-3’), GAPDH (forward 5’-ACCACAGTCCATGCCATCAC-3’ and reverse 5’-TCCACCACCCTGTTGCTGTA-3’), MHCI (forward 5’-CGCACAGAGATACTACAACC-3’ and reverse 5’-ATCTGGGAGAGACAGATCAG -3’)

### Western blotting

BMDC were harvested for lysis after 48 hrs of transfection. Cell suspension in lysis buffer (150mM NaCl, 50mM Tris-Cl pH 8.0, 0.1% SDS, 1% Triton X-100, 0.5% sodium deoxycholate, 50mM DTT, 1mM EDTA, 5% glycerol, 5mM NaF, 2mM Na_3_VO_4_, 50mM PMSF and Protease Inhibitor Cocktail) was incubated on ice for 20 min Following centrifugation at 12,000 rpm for 10 min at 4°C, SDS-PAGE was run with 10ug of protein. After PVDF membrane transfer, blocking was performed with 5%BSA in TBST followed by primary antibody incubation at 4°C overnight. After washing with TBST 3X, bolt was incubated with HRP conjugated secondary antibody for 2 hr at room temperature. Chemiluminescence reagent was used for developing blot in chemidoc. Band intensities were measured by GelQuant software.

### ELISA

Level of secreted Wnt5A by BMDC was measured by coating ELISA plate wells with 100ul of media in which BMDC were cultured. After washing with PBST (0.05% Tween in PBS) wells were blocked with 1% BSA for 2 hr at room temperature, and washed thereafter. Subsequently, after incubation with Wnt5A primary antibody overnight at 4°C, wells were washed again with PBST and incubated with HRP conjugated secondary antibody for 2 hr at room temperature. TMB substrate was used for developing and the reaction was stopped by 250mM HCL and absorbance was measured at 450nm in an ELISA reader.

### Confocal microscopy

For estimation of DQ-OVA intensity, washed and dried poly-l-lysine coated chamber slides were used for culturing BMDC. Following treatment with rWnt5a (100ng/ml) or PBS (vehicle control) for 6 hrs, or 47 hrs post-transfection, BMDC were pulsed with DQ-OVA (5 ug/ml) for 45 min, washed thereafter with cold PBS 3X, fixed in 3% paraformaldehyde for 15 min at room temperature and washed again with PBS 3X. rWnt5a treated BMDC were stained with phalloidin (diluted 1:2000 in PBST with 2.5% BSA) for 15 min, counterstained with DAPI (1:3000) for 10 min, and washed with PBST 3X. Transfected BMDC were incubated with CD11c antibody at room temperature for 1 hr, counterstained with DAPI, and washed with PBS 3X. All treated samples were mounted in 60% glycerol or prolong glass antifade for confocal microscopy [63X magnification (1X or 2X zoom) of Leica TCS-SP8 confocal microscope].

For PA28*α*/β staining, transfection and DQ-OVA pulsing of BMDC were carried out as described above on chamber slides. Subsequently, after fixation in 3% paraformaldehyde, cells were permeabilised for 10 min at room temperature by 0.1% Triton X -1%BSA in PBS, washed with PBS 3X, and blocked for 1 hr at room temperature with 1% BSA in PBST. Treated BMDC were incubated overnight at 4°C separately with either PA28*α* or PA28β primary antibody diluted in PBST with 1% BSA. Next day cells were washed with PBS 3X and incubated with secondary antibody (diluted in 1% BSA) for 2 hr at room temperature. After counterstaining with DAPI for 10 min at room temperature the cells were washed 3X with PBS and mounted in prolong glass antifade for confocal microscopy [63X magnification (2X zoom) of Leica TCS-SP8 confocal microscope].

For estimating T cells attached to BMDC transfected with either Wnt5A siRNA or control, the chamber slide grown cells were incubated with ovalbumin (100ug/ml) about 28 hr post-transfection for the rest of the transfection period. After washing, the OVA pulsed transfected BMDC were co-cultured with MAC purified CD8 T cells from ovalbumin immunized mice for 1 hr at 37°C. After washing with PBS 3X to remove unbound T cells samples were fixed in 3% paraformaldehyde for 15 min at room temperature. BMDC-T cell conjugate was analyzed by confocal micrscopy. In some experiments, anti-CD11c and anti-CD8 staining were performed for identifying BMDC and CD8 T cells. Samples were mounted in 60% glycerol for visualization of BMDC-T cell conjugates by confocal microscopy [63X magnification (3X or 2X zoom) of Leica TCS-SP8 confocal microscope].

### Flow cytometry

For estimation of antigen uptake and antigen processing, transfected BMDC (47 hr post-transfection) were pulsed with either FITC-Ova (5ug/ml) or DQ-OVA (5ug/ml) for 45 min, after which cells were washed with cold PBS and stained for 1 hr with anti-CD11c antibody for subsequent FACS (BD.LSR Fortessa Cell analyzer).

For estimation of antigen presentation, transfected BMDC, either stimulated with OVA (100ug/ml) or left unstimulated were harvested, washed with PBS with 0.5% BSA, and stained with anti-CD11c, anti-MHCI (H-2K[d]), and anti-MHCII (I-A/I-E) for 1 hr at 4°C. After washing cells were analyzed by FACS.

For analysis of CFSE dye dilution and proliferation of OVA-sensitized CD8 T cells in coculture with BMDC, CD8 T cells were purified from total spleen and lymph node of OVA immunized mice by MACs column mediated negative selection after RBC lysis (“Red Blood Cell Lysis Buffer,” 2006), and thereafter labeled with 2uM CFSE in PBS for 10 min at 37°C (https://flowcytometry.utoronto.ca/wp-content/uploads/2016/01/Proliferation-CFSE.pdf), before co-culturing with BMDC. CD8 T cell purity and CFSE labeling were confirmed by FACS, before setting up the co-culture. After 4 days of co-culture, floating and loosely adherent cells were harvested, stained with anti-CD3 and anti-CD8 antibodies for 1 hr at 4°C, for identifying CD8 T cells and PI was added before FACS acquisition for excluding dead cells. For measuring OVA recall response in Wnt5A+/+ and Wnt5A+/-mice by CFSE dye dilution, cells harvested from mice spleens and lymph nodes were labeled with CFSE after RBC lysis. Following incubation with OVA (100ug/ml) for 4 days, cells were harvested and stained with anti-CD3 and anti-CD8 antibody for 1 hr at 4°C. In a subset of similar experiments, MHCI tetramer staining was performed the same way before CD8 and CD3 staining. For all samples either PI or 7AAD was used as cell viability dye before FACS acquisition.

Intracellular cytokines were measured after 48 hrs of culture following Brefeldin (5ug/ml) treatment for the last 4 hrs. Harvested cells were fixed with 1% paraformaldehyde for 15 min at room temperature, permeabilized with 0.1% Tween-20-1%BSA-PBS for 10 min at room temperature, washed with PBST 3X, and finally stained with anti-CD3, anti-CD8, anti-IL2, anti-IFNY, and anti-GranzymeB antibodies in PBST (PBS + 0.05%Tween) with 1% BSA, for 1 hr at 4°C, before FACS acquisition.

### Statistical analysis

Statistical analysis was done using Student’s paired t-test for densitometry units, MFI of DQ-OVA (FACS), MFI of MHCI, MFI of MHCII, proliferative index of Wsi vs Csi, ELISA, and for adherent T cells per 100 DC. Student’s unpaired t-test was used for DQ-OVA intensity measurement (confocal), % overlap (confocal), PA28*α* and PA28β intensity (confocal), proliferative index of wild type vs heterozygous mice, cytokines of wild type vs heterozygous mice, and for CD44^high^CD62L^low^ %.

## Results

### Wnt5A promotes processing of internalized antigen in association with actin assembly

Given the involvement of Wnt5A signaling in cytoskeletal actin dynamics, a major player in antigen uptake (Roper et al., 2019), we investigated how Wnt5A signaling influences antigen processing, this being a prerequisite for antigen presentation and T cell activation. To this end we used mouse bone marrow derived dendritic cells (BMDC) as model antigen processing/presenting cells and ovalbumin (OVA) as a model foreign antigen.

BMDC was generated from the bone marrow of BALB/c mice following published protocols (P. Matheu et al., 2008), and purity was confirmed by flow cytometry using CD11C antibody (Figure S1A). Subsequently, we demonstrated that BMDC express both the secreted ligand Wnt5A as well as Frizzled5, an established receptor for Wnt5A (Blumenthal et al., 2006; Maiti et al., 2012; Naskar et al., 2014), indicating that BMDC are capable of both paracrine and autocrine Wnt5A signaling (Figure S1B). In order to evaluate the influence of Wnt5A signaling on OVA processing, we treated BMDC culture with BODIPY conjugated DQ-Ovalbumin (DQ-OVA), which emits bright green fluorescence while being proteolytically processed (Olatunde et al., 2018; Rivera-Gil et al., 2009). The extent of antigen processing, in the form of DQ-OVA cleavage was evaluated by the intensity of fluorescence emitted.

We demonstrated increased association of assembled actin with DQ-OVA processing in recombinant Wnt5A (rWnt5A) treated BMDC as compared to the corresponding PBS control. Figure 1 (Panel Ai) is a confocal microscopic representation of higher intensities of both phalloidin-red (measure of filamentous actin) and DQ-OVA-green, and higher percentage of DQ-OVA-phalloidin co-localization (yellow) in rWnt5A treated BMDC than the vehicle (PBS) treated control. Panels Aii and Aiii denote the level of rWnt5A-induced increase in DQ-OVA intensity and DQ-OVA-phalloidin colocalization. This observation clearly indicates that the Wnt5A-Actin axis, which has been described explicitly in previous publications (Chakraborty et al., 2017; Jati et al., 2018) promotes antigen processing. To further assess the dependence of antigen processing on Wnt5A we examined the effect of siRNA mediated silencing of Wnt5A in BMDC on DQ-OVA processing. Both Wnt5A siRNA transfected and scramble siRNA (control) transfected BMDC were treated with DQ-OVA for about 45 min to enable OVA internalization and processing, following which intracellular fluorescence was measured in each group of cells. The decision to treat BMDC with DQ-OVA for 45 mins was based on prior experiments where BMDC were treated with DQ-OVA under different conditions (Figure S2A). Confocal microscopy revealed significantly lower DQ-OVA fluorescence in the Wnt5A siRNA sets than in the corresponding control sets (Figure 1: Panels Bi and Bii). Panels B iii-v demonstrate the siRNA-mediated reduction in Wnt5A expression in BMDC, both at mRNA and protein level. Flow cytometry, separately conducted on DQ-OVA treated BMDC transfected with either Wnt5A siRNA or control siRNA yielded results that were very similar to those yielded by confocal microscopy (Figure 1: Panels Ci and Cii) corroborating that Wnt5A signaling is required for antigen processing. There was no significant influence of Wnt5A on OVA uptake (Figure S3), indicating that the observed effect on OVA processing was not due to any notable alteration in OVA uptake.

**Figure 1.**
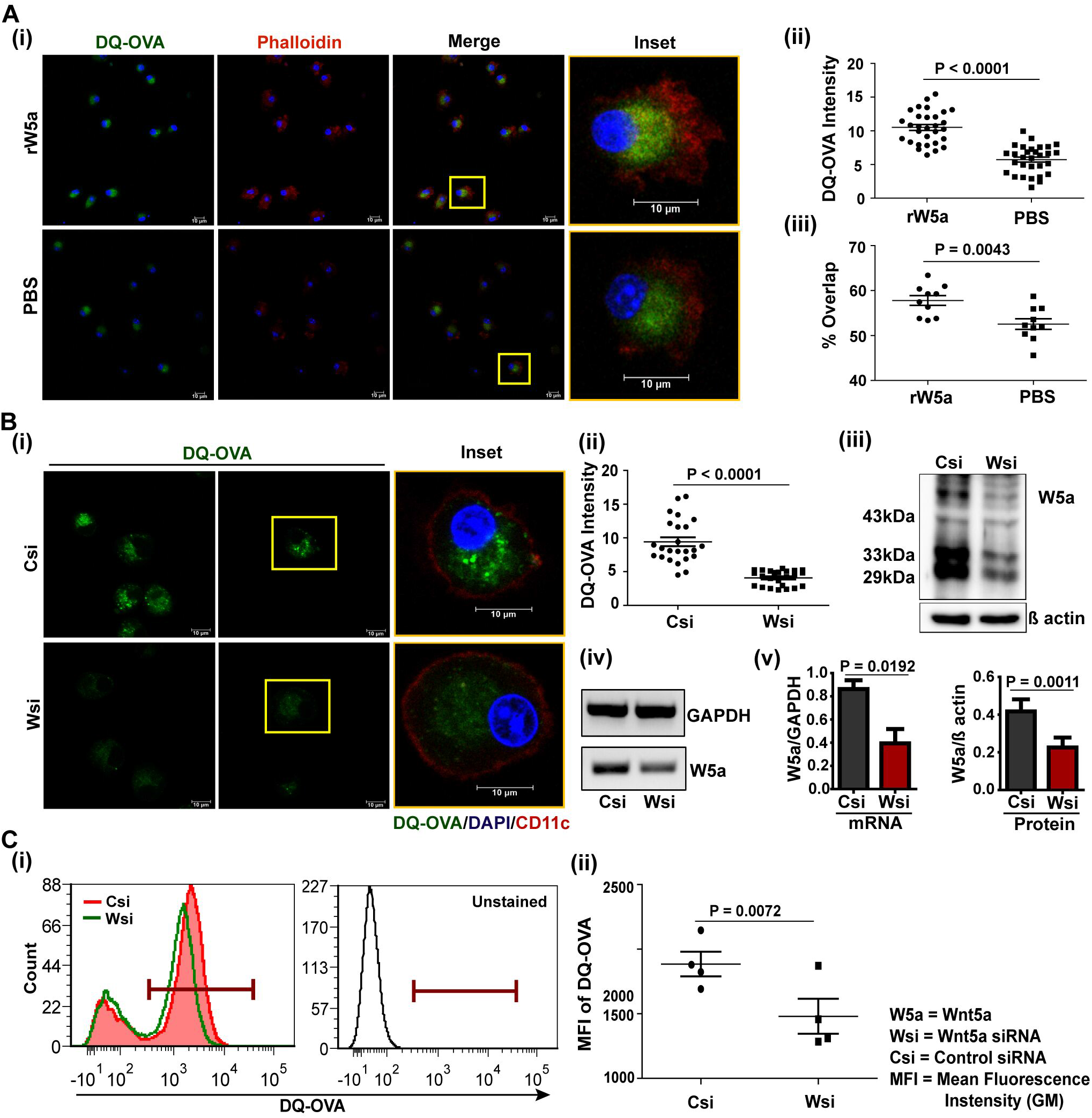
Wnt5A promotes DQ-OVA processing: A: rWnt5A mediated increase in phalloidin (i) and DQ-OVA (i and ii), as demonstrated by confocal microscopy. (iii) rWnt5A mediated increase in DQ-OVA and phalloidin colocalization. B: Similar experiments (i and ii) showing decrease in DQ-OVA intensity by siRNA mediated depletion of Wnt5A. Wnt5A depletion is shown at both protein and mRNA levels (iii, iv, v). C: Flow cytometry demonstrating a decrease in DQ-OVA fluorescence in Wnt5A depleted cells as compared to control. A, B, and C: For intensity measurement (n = 2), each dot represents a cell. For colocalization measurement (Manders overlap coefficient, n = 2), each dot is a field. For confocal microscopy, magnification was 63X (1X or 2X zoom). For flow cytometry (n = 4) FITC (marker) gate was selected based on unstained. Data represent mean ± SEM.

### Wnt5A facilitated antigen processing is proteasome-mediated

The fact that DQ-OVA processing, both with and without exogenous Wnt5A is to a large extent mediated by MG132 inhibitable proteasome activity (Figures S4A and S4B), raised the question whether Wnt5A facilitated DQ-OVA processing is in fact linked with proteasome function. Accordingly, in view of the demonstrated involvement of the proteasome regulator PA28*α*-PA28β hetero-oligomer in antigen processing/presentation (Rechsteiner et al., 2000; van Hall T et al., 2000), we examined if PA28*α* and PA28β are associated with Wnt5A assisted antigen processing. Initially, we demonstrated colocalization of fluorescing DQ-OVA and PA28β in BMDC (Figure S4C), which is in compliance with the observed blockade in DQ-OVA processing by the proteasomal inhibitor MG132 (Bao et al., 2016; Sakabe et al., 2012). Subsequently, we demonstrated significantly reduced colocalization of PA28*α* and PA28β with DQ-OVA in BMDC sets where Wnt5A was depleted by Wnt5A siRNA (Wsi), in comparison to the control sets (Csi), by confocal microscopy (Figure 2: Panels Ai, Aii & Ci, Ciii). Wnt5A depletion in fact also led to a significant reduction in the levels of both PA28*α* and PA28β (Panels Ai, Aii & Cii, Civ). These results indicate that Wnt5A signaling regulates both the expression of the PA28 proteasome regulator subunits as well as their association with the target antigen being processed. We furthermore examined the effect of siRNA-mediated Wnt5A depletion in BMDC on PA28*α*-PA28β co-localization by confocal microscopy. As demonstrated in Figure 2, Panels B & C v, PA28*α*-PA28β co-localization was significantly reduced in Wnt5A depleted BMDC sets as compared to the corresponding controls, implying that Wnt5A signaling regulates PA28*α*-PA28β assembly. The extent of colocalization (PA28*α*/PA28β-DQ-OVA, PA28*α*-PA28β) was measured using Manders overlap coefficient (Dunn et al., 2011). Similar results were obtained using Pearson’s coefficient (Adler and Parmryd, 2010).

**Figure 2.**
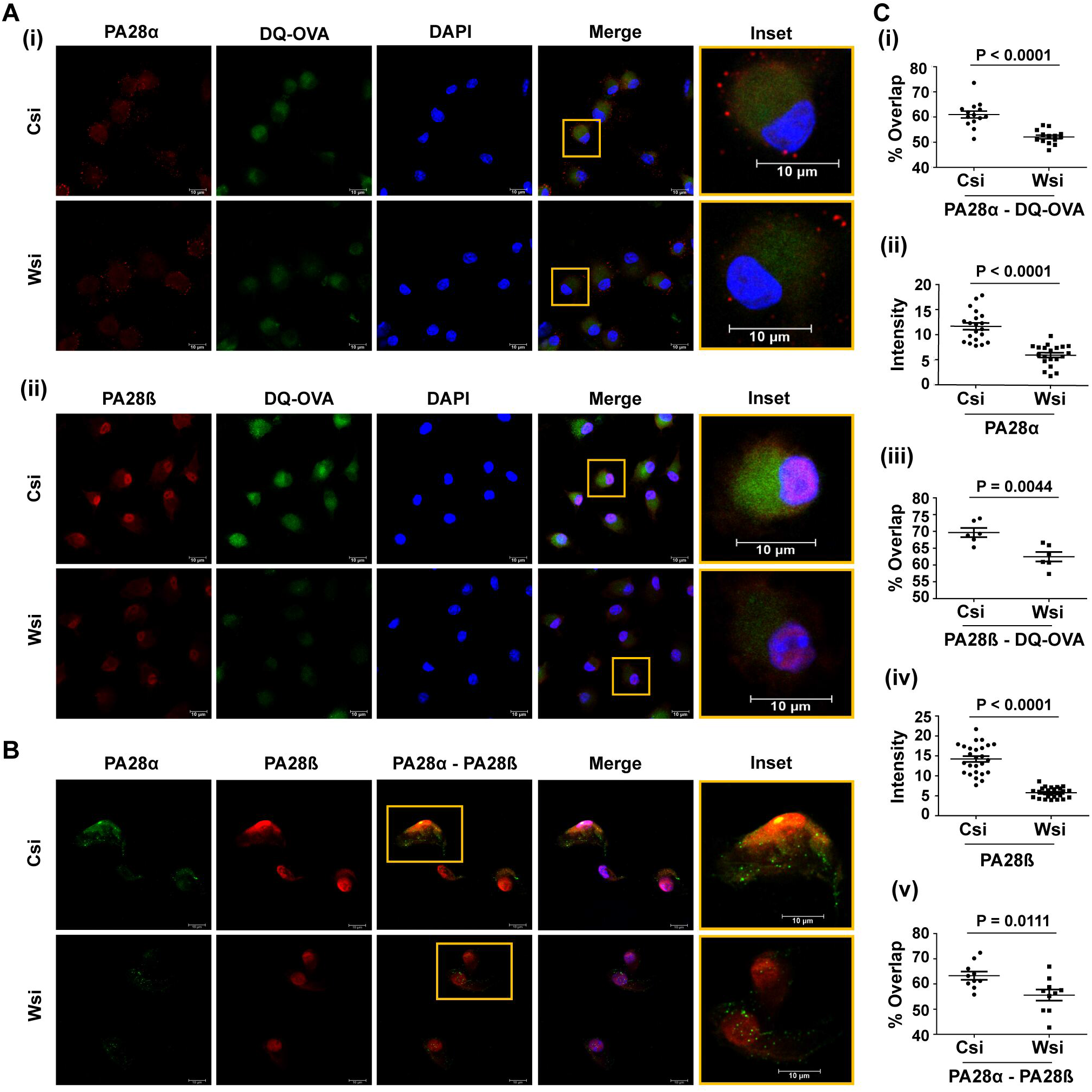
Wnt5A mediated DQ-OVA processing is associated with PA28*α*-PA28β: A and C: Confocal microscopy demonstrating decrease in DQ-OVA-PA28*α* and DQ-OVA-PA28β overlap (A, Ci, Cii) in Wnt5A depleted (Wsi) BMDC as compared to control (Csi). B and C (v): Decrease in the association between PA28*α* and PA28β by Wnt5a depletion, as measured by overlap intensity (Manders overlap coefficient, n = 2). Magnification 63X (2X zoom). Data represent mean ± SEM.

Taken together, our experimental results reveal that Wnt5A signaling promotes antigen processing by acting in conjunction with both actin and proteasome dynamics. Accordingly, it was important to examine if Wnt5A facilitated antigen processing also influences antigen presentation and CD8 T cell activation.

### Wnt5A mediated antigen processing correlates with cross presentation to MHC Class I and antigen specific CD8 T cell activation

In order to examine if Wnt5A-proteasome activity correlates with antigen cross-presentation to MHC Class I and CD8 T cell activation, an outcome of proteasome-mediated antigen processing (Rechsteiner et al., 2000), we used BMDC, OVA and OVA primed CD8 T cells.

First, we demonstrated that OVA treatment of BMDC, which involves processing of the internalized OVA leads to an increase in the expression of surface MHC class I molecules (Figure S5A), thus validating antigen cross-presentation in our system. Subsequently, we addressed if OVA-dependent MHC Class I expression is in fact influenced by Wnt5A signaling. To this end, surface MHC molecule expression in Wnt5A depleted (Wnt5A siRNA transfected) BMDC was compared to that in the control (scramble siRNA transfected) by flow cytometry both with and without OVA pulsing. As depicted in Figure 3 (Panels A and Ci), estimation of the mean fluorescence intensity (MFI: geometric mean) of surface MHC revealed significant inhibition in MHC Class I surface expression after OVA stimulation in Wnt5A depleted condition, when compared to the control. The slight decrease in surface MHC Class I expression in the Wnt5A siRNA sets compared to the corresponding controls even in the absence of OVA stimulation (displayed in the same panels) could be due to decreased recycling of MHC class I to the plasma membrane (Montealegre and van Endert, 2019). There was no detectable change in the level of MHC Class I mRNA expression in BMDC upon depletion of Wnt5A both in OVA stimulated and un-stimulated states (Figure 3, Panel Cii), indicating that Wnt5A dependent MHC Class I surface expression was not due to induction of MHC Class I gene expression. Wnt5A depletion in BMDC did not affect the surface expression of MHC Class II either in presence or absence of OVA stimulation significantly (Figure 3, Panel B and Ciii). Thus the effect of Wnt5A signaling dependent antigen processing was mostly on the proteasome controlled antigen cross-presentation by MHC Class I and not on the usually endosome controlled antigen presentation by MHC Class II (Heath and Carbone, 2001; Joffre et al., 2012). This finding was in agreement with the observed association of Wnt5A signaling with the proteasome regulatory complex subunits (Figure 2), which are involved in antigen presentation by MHC Class I (Sijts et al., 2002).

**Figure 3.**
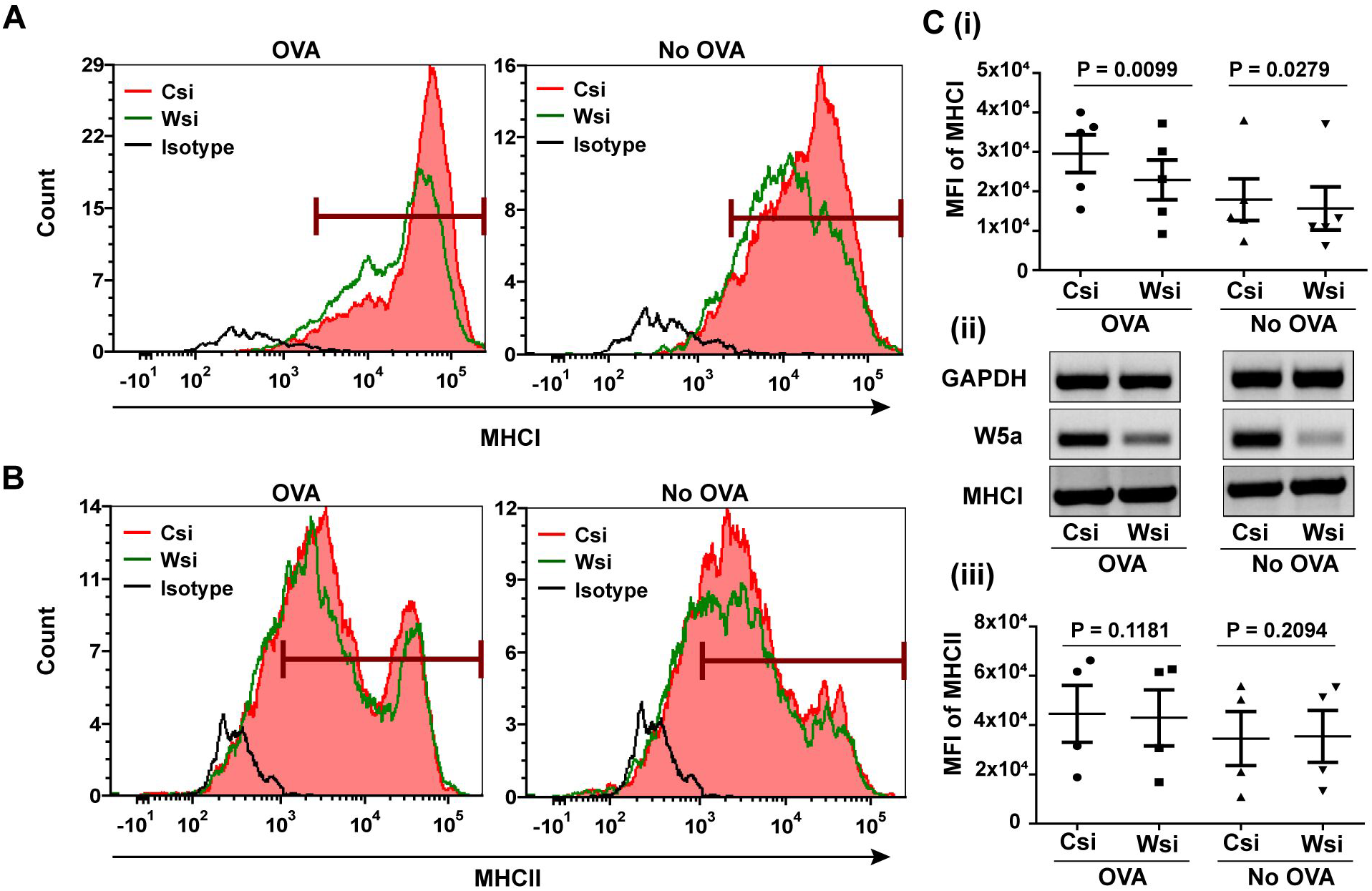
Wnt5A promotes antigen cross-presentation: A and C: FACS histogram (A) and corresponding MFI plot (C) of surface MHCI (PE) on BMDC with and without OVA pulsing, in Wnt5A depleted condition (Wsi) as compared to control (Csi) (n = 5) MHCI gene expression is not affected by Wnt5A depletion (Cii). B and C (iii): FACS demonstrating no significant effect of Wnt5A depletion on MHC ClassII (Per CP-Cy 5.5) surface expression. Marker gate based on isotype control (n = 4). Data represent mean ± SEM.

Wnt5A facilitated antigen cross-presentation on MHC Class I was furthermore validated by the proliferation of OVA-sensitized CD8 T cells while in co-culture with OVA pulsed BMDC. To this end, OVA-sensitized CD8 T cells harvested from the spleen and lymph node of OVA immunized BALB/c mice (explained in Materials and Methods) were labeled with CFSE and placed in culture with BALB/c derived OVA pulsed BMDC, which were transfected with either Wnt5A siRNA or scramble siRNA (control) before OVA pulsing. Additionally, a portion of the BMDC, either transfected with Wnt5A siRNA or scramble siRNA, but not stimulated with OVA was placed in similar co-culture with CD8 T cells to serve as “no OVA” un-stimulated reference for each “Wnt5A siRNA OVA” set or “scramble siRNA OVA” set. After 4 days of co-culture cells were harvested and CD8 T cell proliferation was estimated by flow cytometry from the CFSE distribution therein. For both, Wnt5A siRNA (Wsi)-BMDC/T cell co-culture, and control/scramble (Csi)-BMDC/T cell co-culture, cells were gated based on the expression of CD3 and CD8 after live-dead cell gating with propidium iodide (PI), as represented by examples in Figure 4, Panel A, and the proliferation for “OVA stimulation” compared to “no OVA” was calculated from the CFSE intensity histogram using FCS Express software as represented by examples in Figure 4, Panel B. For each “OVA sample”, OVA-specific proliferation was estimated based on the median of the corresponding “no-OVA” sample as un-stimulated control and the median for unstained cells as a reference for auto-fluorescence. We noted that depletion of Wnt5A expression in BMDC by siRNA resulted in significantly reduced OVA-specific CD8 T cell proliferation as demonstrated by the values of proliferative index in Figure 4, Panel C. Wnt5A depletion also led to a considerable reduction in the number of CD8 T cells in conjugation with BMDC about 1 hr after co-culture, implying subdued CD8 T cell recognition by the BMDC plausibly on account of impaired OVA processing and presentation (Figure 4, Panel D: i & ii). Association between BMDC (CD11c: red) and CD8 T cell (CD8: green) was validated separately by confocal microscopy (4D: iii). Impaired OVA-specific CD8 T cell activation by siRNA mediated Wnt5A depletion in BMDC after OVA stimulation was reflected in their reduced levels of intracellular IFN*γ*, IL2, and granzyme B (GRB) as depicted in Figure 4, Panel E (Ei represent specific examples of OVA-specific IFN*γ*, IL2 and GRB response by FACS, and Eii represent the overall decrease in CD8 T cell activation upon Wnt5A depletion). Integrated Mean Fluorescence Intensity (iMFI: % positive events X geometric mean) was used for projection of T cell activation because both % positive events, as well as geometric mean, account for activation (Darrah et al., 2007). The robustness of Wnt5A responsive CD8 T cell response after OVA pulsing was much reduced when the CD8 T cells were harvested from OVA unimmunized mice (Figure S5B), validating the efficacy of OVA immunization for the generation of OVA-sensitized CD8 T cells. The purity of isolated CD8 T cells is depicted in Figure S5C.

**Figure 4.**
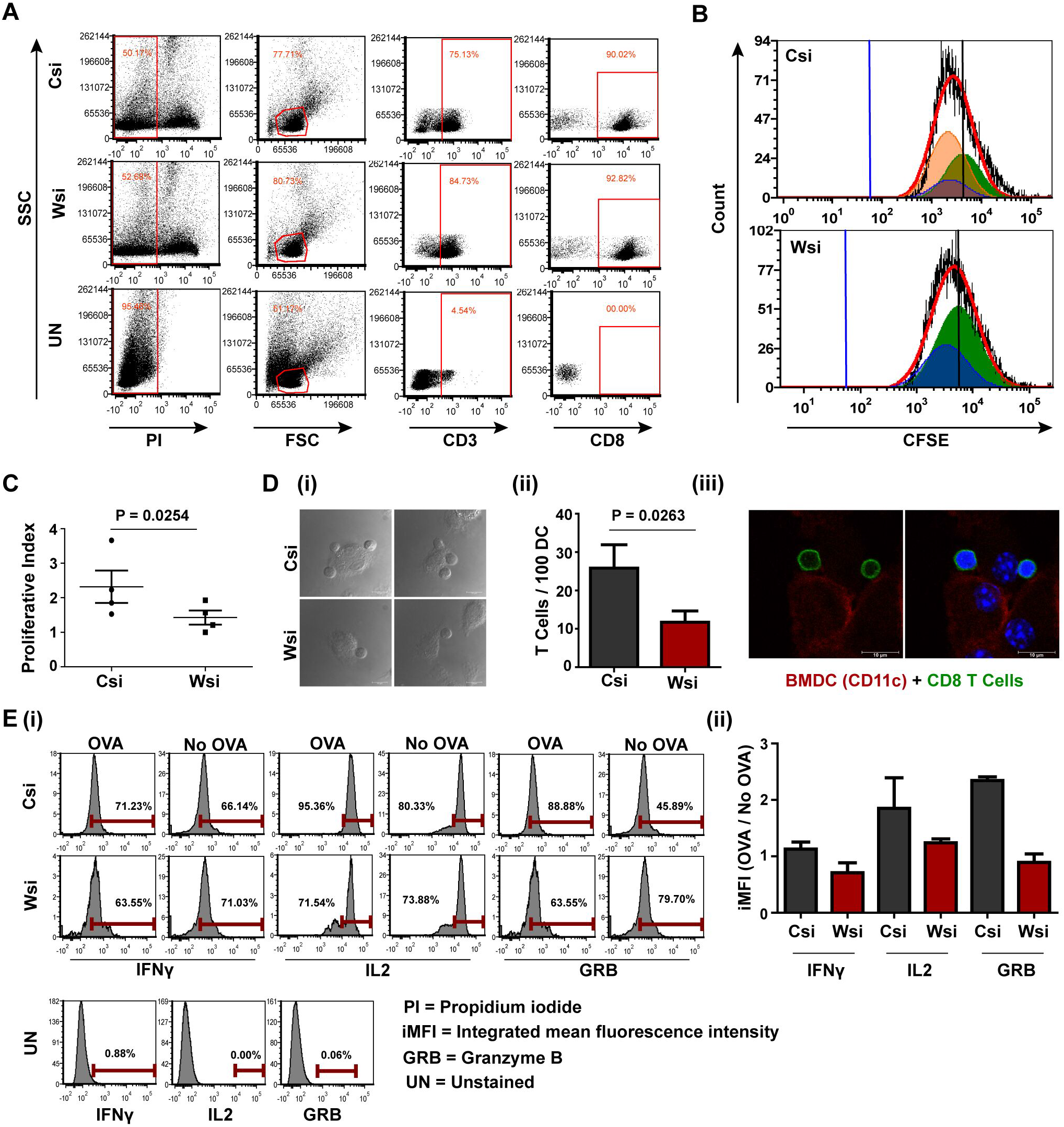
Wnt5A mediated OVA processing in BMDC promotes CD8 T cell activation: A and B: Representation of FACS gating of OVA responsive CD8 T cells (A) harvested from OVA pulsed BMDC (Wsi vs Csi) -T cell co-culture and their CFSE (FITC) histogram plot (B). C: CD8 T cells in co-culture with Wnt5A depleted OVA pulsed BMDC have less proliferation index than the corresponding control (Csi) (n = 4). D: CD8 T cells adhere less to Wnt5A depleted BMDC (n = 3). Magnification 63X (3X or 2X zoom). E: Representative examples of intracellular IL2 (APC-R700), IFN*γ* (PE-CF594), and GRB (PE-Cyanine 7) in CD8 T cells in co-culture with OVA pulsed BMDC (Wsi vs Csi) as analyzed by flow cytometry (n = 2). Data represent mean ± SEM.

As a whole, these results indicate that Wnt5A signaling supports antigen cross-presentation and CD8 T cell activation. On account of the negligible influence of Wnt5A on MHC II surface expression in response to antigen stimulus in our system (Figure 3, Panels B and C), we did not analyze the effect of Wnt5A on CD4 T cell activation from the angle of antigen presentation.

### Antigen specific CD8 T cell recall response to OVA antigen is impaired in Wnt5A heterozygous (WntA+/-) background

Having demonstrated the requirement of Wnt5A signaling in antigen cross-presentation and concomitant CD8 T cell activation, it was important to investigate if Wnt5A signaling influences the potential for CD8 T cell recall response to antigen following immunization. Accordingly, we compared the extent of OVA-specific CD8 T cell activation and recall response in Wnt5A deficient (Wnt5A +/-) mice with that of the wild type (Wnt5A +/+) counterparts after immunization with OVA antigen. The OVA immunization protocol and FACS gating strategy for assessing CD8 T cell activation and recall response are outlined in Materials and Methods and Figure 5 Panel A, respectively. In essence, following OVA immunization with two doses of antigen, both sets of mice, Wnt5A wild type and Wnt5A heterozygous were sacrificed, and CD8 T cell recall response to antigen was evaluated by flow cytometry after stimulating splenocyte cultures derived from each set with fresh OVA antigen. For each OVA pulsed splenocytes/lymph node culture, a similar “no OVA” culture was used as reference. CD8 T cell proliferation was measured after 4 days, through analysis of CFSE proliferation histogram on live CD3^**+**^CD8^**+**^ gated cells in the same was as described before in Figure 4. Here, 7AAD was used instead of PI for live/dead cell gating. We observed significantly lower CD8 T cell proliferation in response to OVA stimulation in the splenocyte/lymph node cultures of the Wnt5A heterozygous mice as compared to those of the corresponding wild type cohorts (Figure 5, Panels B and C), indicating that OVA-specific CD8 T cell recall response after immunization is compromised in the Wnt5A heterozygous background. To further ensure the specificity of CD8 T cell response during immunization and *in vitro* recall response to antigen, we used labeled SIINFEKL bound MHCI tetramer as a tool for identification of OVA-specific CD8 T cells in a subset of the experiments represented in Figure 5B. We calculated the number of SIINFEKL-MHCI tetramer bound CD8 T cells from the proliferated (CFSE less) population (detailed procedure explained in Materials and Methods). We noted that the number of proliferating MHCI tetramer bound CD8 T cells was relatively less in the Wnt5A heterozygous mice as compared to the controls (Figure 5, Panel Di). Furthermore, the percentage of CD44^high^CD62L^low^ phenotype among the proliferating MHC tetramer binding CD8 T cells was also low in the Wnt5A heterozygous mice as compared to the wild type, as demonstrated by density plot (Figure 5, Panels Dii and Diii), indicating compromised differentiation to an effector memory phenotype during the *in vitro* recall response to antigen (Honey et al., 1999; Roberts and Woodland, 2004; Zhai et al., 2003), perhaps on account of inefficient antigen priming during immunization.

**Figure 5.**
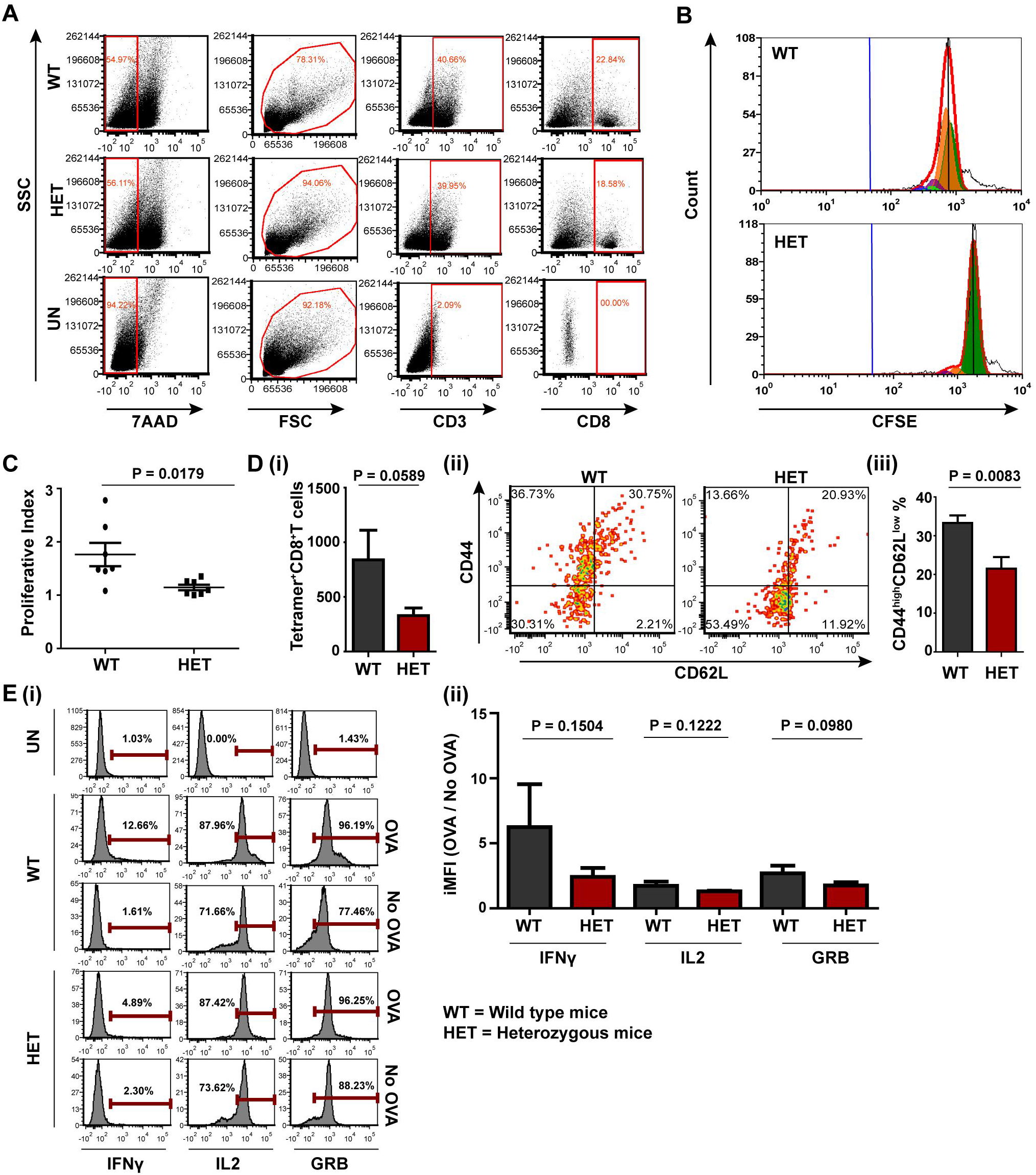
CD8 T cell recall response of OVA: A and B: Representative examples of OVA-specific CD8 T cell gating by FACS (A) and their CFSE histogram (B). C: CD8 T cells from heterozygous mice have shown less PI than those of control (Wild type mice) in response to OVA stimulation. D: Higher number of MHCI-tetramer (PE) binding CD8 T cells in WT, than those of Heterozygous (i) with more pronounced effector phenotype (ii and iii) CD44^high^ (APC) CD62L^low^(PE-CY7). Quadrant is based on secondary control of CD62L. E (i and ii): Representative examples of intracellular IL2 (APC-R700), IFN*γ* (PE-CF594), and GRB (PE-Cyanine 7) in CD8 T cells from Wild type and Heterozygous mice. Data represent mean ± SEM.

Increased OVA-specific CD8 T cell proliferation in the wild type splenocyte/lymph node cultures correlated with augmented levels of OVA-specific IFN*γ*, IL2, and GRB in a similar set of flow cytometry experiments (Figure 5, Panel E: i and ii), as described in Figure 4, implying an activated state of CD8 T cells with potential toxicity (Mahajan et al., 2003; Yoshida et al., 2006). Overall, these results indicate that OVA antigen-specific CD8 T cell priming is markedly better in the Wnt5A wild-type mice than their heterozygous counterparts. The lack of appropriate CD8 T cell priming in the Wnt5A heterozygous mice could very well be an outcome of ineffective processing of antigen and its cross-presentation to CD8 T cells.

## Discussion

Cell surface presentation of antigenic epitopes of infectious agents on the MHC Class I molecules of antigen-presenting cells is associated with CD8 T cell activation, a vital component of cell-mediated immunity. Dearth of expansion of antigen-specific CD8 T cells, which serve to build CD8 T cell memory to exposed antigens, leads to impairment in cell-mediated immunity and may cause susceptibility to infections (Huang et al., 2014; Velilla et al., 2006). Although there has been much research performed and extensive knowledge gained along these lines, several questions remain in relation to how antigen-presenting cells process and present internalized foreign antigens toward the generation of antigen responsive CD8 T cells for future recall response to antigen.

In this study, we evaluated the role of Wnt5A signaling in antigen processing, cross-presentation, and CD8 T cell activation in light of its documented involvement in cytoskeletal actin remodeling, a core element of intracellular vesicle movements and antigen presentation (Huang et al., 2011; Schuh, 2011; Semenova et al., 2008). In order to precisely focus on host factors that direct antigen processing/presentation toward T cell activation, we did not apply any co-stimulation, such as LPS, in addition to antigen pulsing in our system.

Using mouse BMDC as antigen-presenting cells and OVA as a model antigen we demonstrated that Wnt5A promotes antigen cross-presentation and CD8 T cell activation by regulating actin and PA28 *α*/β dynamics. (Figures 1-4). Although not demonstrated in this study, the observed influence of Wnt5A on PA28 *α*/β expression could be mediated through IFN*γ*, which has been previously found to be regulated by Wnt5A signaling (Naskar et al., 2014; Valencia et al., 2014). In light of the association of proteasome activity with actin dynamics (Haarer et al., 2011), Wnt5A directed effects on actin and proteasome regulation in the context of antigen processing/presentation may very well be coordinated.

Impairment in the activation of antigen-responsive CD8 T cells was observed in a Wnt5A depleted background in Wnt5A heterozygous (Wnt5A+/-) mice as illustrated by their significantly reduced CD8 T cell recall response to OVA antigen in comparison to the wild type cohorts following immunization with OVA. Inhibited cross-presentation of OVA to CD8 T cells in the heterozygous mice was evident from the reduced numbers of SIINFEKL-MHCI tetramer binding activated CD8 T cells in the recall response. On the whole, these results suggest that host Wnt5A signaling supports antigen priming of CD8 T cells through antigen processing and cross-presentation, and as a corollary, facilitates the generation of antigen-specific memory that is reflected in the form of recall response to antigen (Figure 5).

The outcome of Wnt signaling on T cell response is to a large extent dependent on pathophysiology and the cells under consideration. Wnt5A has been reported to suppress the adaptive immune response of CD8 T cells in melanoma through metabolic programming (Zhao et al., 2018). Wnt3A and Wnt1 also appear to have similar effects on adaptive immunity in different tumor microenvironments (Kerdidani et al., 2019, p. 1; Pacella et al., 2018, p. 1). The level of Tcf-β-catenin mediated transcriptional activation in T cells, on the other hand has been reported to control the fate of T cell memory. (Tiemessen et al., 2014; Utzschneider et al., 2016)

We have studied regulation of CD8 T cell activation and antigen recall response in the context of antigen processing/presentation in relation to actin and proteasome dynamics. Our study suggests that Wnt5A signaling in antigen-presenting cells acts in coordination with actin and proteasome dynamics to aid in antigen-specific CD8 T cell activation for future CD8 T cell recall response to antigen. Identification of host molecules such as Wnt5A that have significant involvement in enrichment of antigen responsive CD8 T cell expansion after immunization may prove useful in the design of effective vaccines.

## Ethics Statement

Animal Ethics Committee of CSIR-IICB (CSIR-IICB-AEC) approved all animal studies in its meeting held on 19^th^ September 2019.

## Author Contribution

MS designed research, analyzed data, and wrote the paper. TRS performed research, contributed to research design, analyzed data, and assisted in writing the paper.

## Funding

This work was supported by the Department of Biotechnology, Govt. of India and institutional funding. TRS was supported by CSIR fellowship.

## Acknowledgements

The authors thank Tanmoy Dalui for FACS, Shounak Bhattacharya for confocal microscopy, CSIR-IICB central instrument facility for instrument support, CSIR-IICB animal house facility for animal breeding & maintenance, Shreyasi Maity for helping in animal handling, Sohum Sengupta, Ananya Ganguly, Deepesh Kumar Padhan for general lab support and Indrajit Sikder for technical assistance.

## Notes

### Competing Interest Statement

The authors have declared no competing interest.

