## supplementary figures for "Wnt5A supports antigen cross-presentation and CD8 T cell activation"

### Supplementary fig: 1

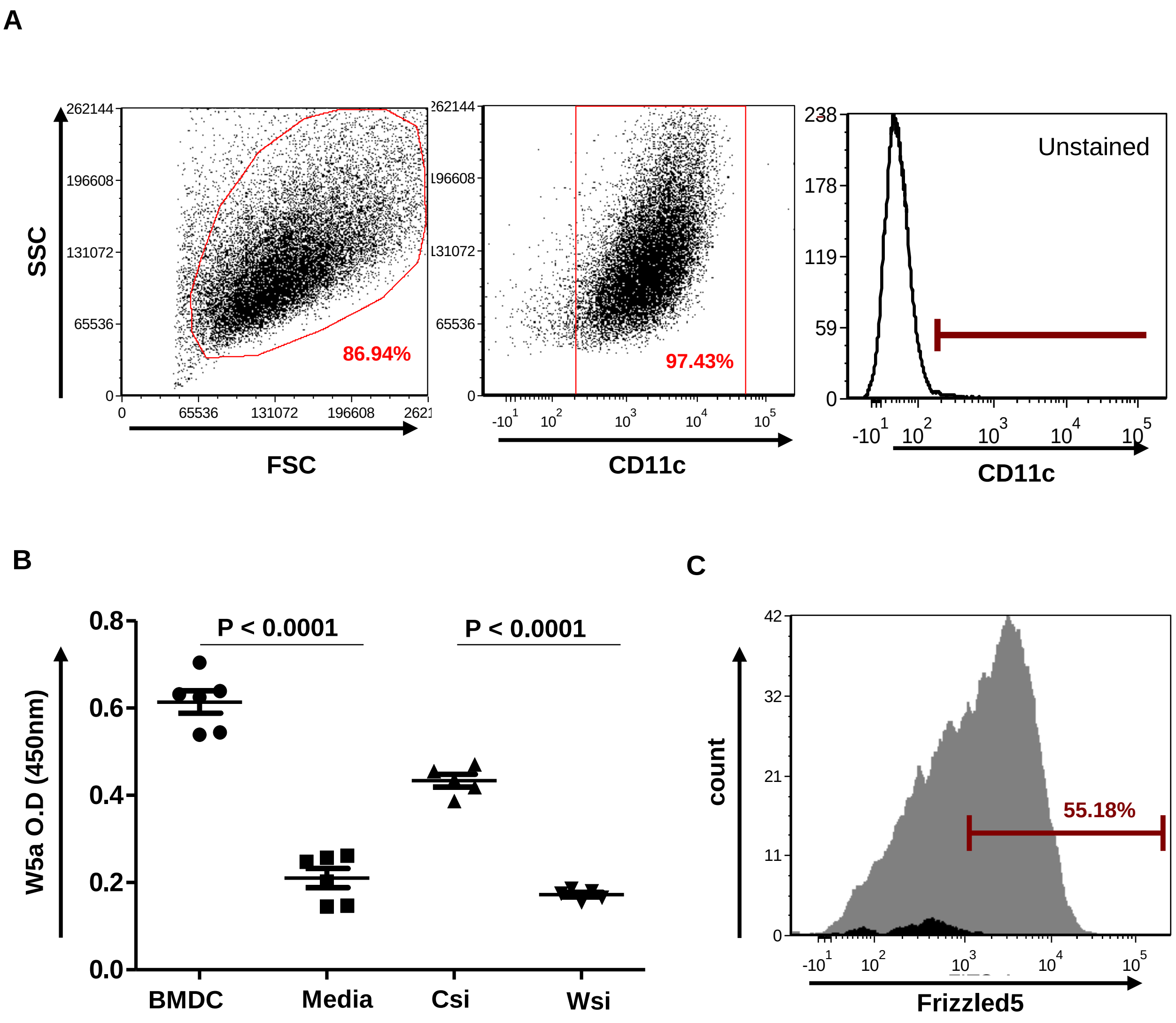


**Figure S1:** **<u>Bone marrow-derived dendritic cells (BMDC) express Wnt5A and Frizzled5 (Wnt5A receptor)</u>**: **A:** Flow cytometry (FACS) of BMDC generated from bone marrow of BALB/c mice demonstrating its purity based estimation of % positive cells expressing CD11c (dendritic cell marker). **B:** ELISA demonstrating secretion of Wnt5A from BMDC as corroborated by its reduction through Wnt5A siRNA mediated depletion of Wnt5A. Cell culture medium without BMDC, and scramble siRNA (control) transfected cells are used as reference for untransfected BMDC and Wnt5A siRNA transfected BMDC respectively. **C:** FACS demonstrating expression of Frizzled5 by BMDC. Isotype control (dark histogram) is the reference

### Supplementary fig: 2

A

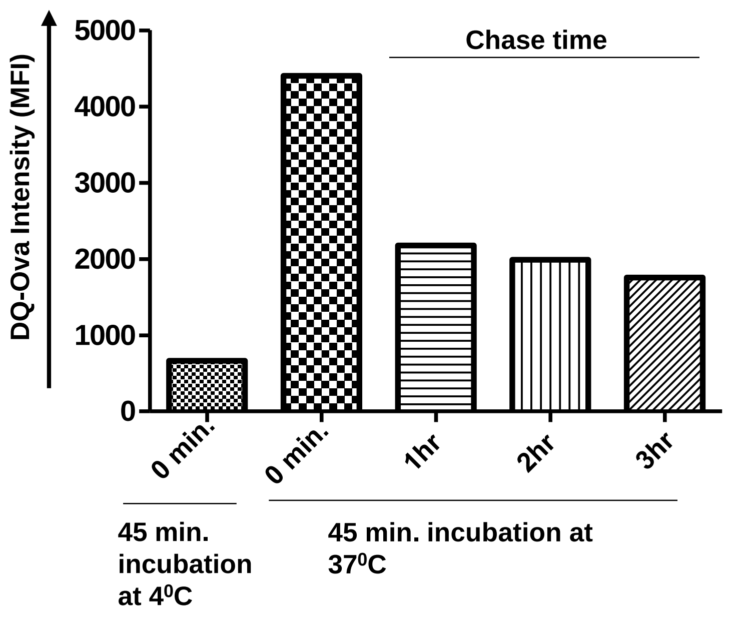


**Figure S2:** **<u>Processing of OVA by BMDC is indicated by DQ-OVA fluorescence</u>:** Incubation of BMDC with DQ-OVA for 45 min at 37$^0$C causes considerable increase in DQ-OVA fluorescence, and 4$^0$C incubation used as a negative control. Fluorescence wanes with chase time. Fluorescence was detected by FACS. MFI: Mean Fluorescence Intensity (Geometric Mean).

### Supplementary fig: 3

A

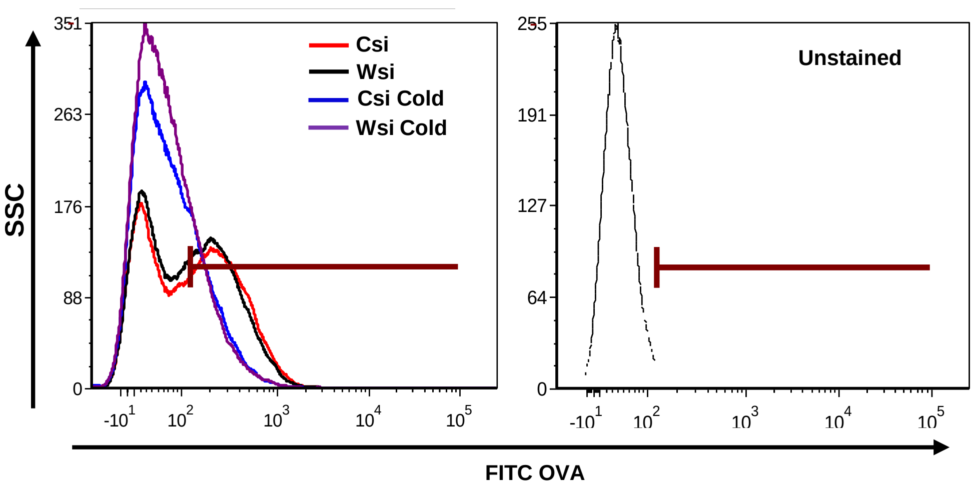


**Figure S3:** **Wnt5A does not influence uptake of OVA:** FACS histogram demonstrating that there is no significant difference in FITC-OVA fluorescence between BMDC transfected separately by Wnt5A siRNA (Wsi) and Control siRNA (Csi) both at cold temperature and $37^{0}$C. FACS gating was based on unstained cells.

Supplementary fig: 4

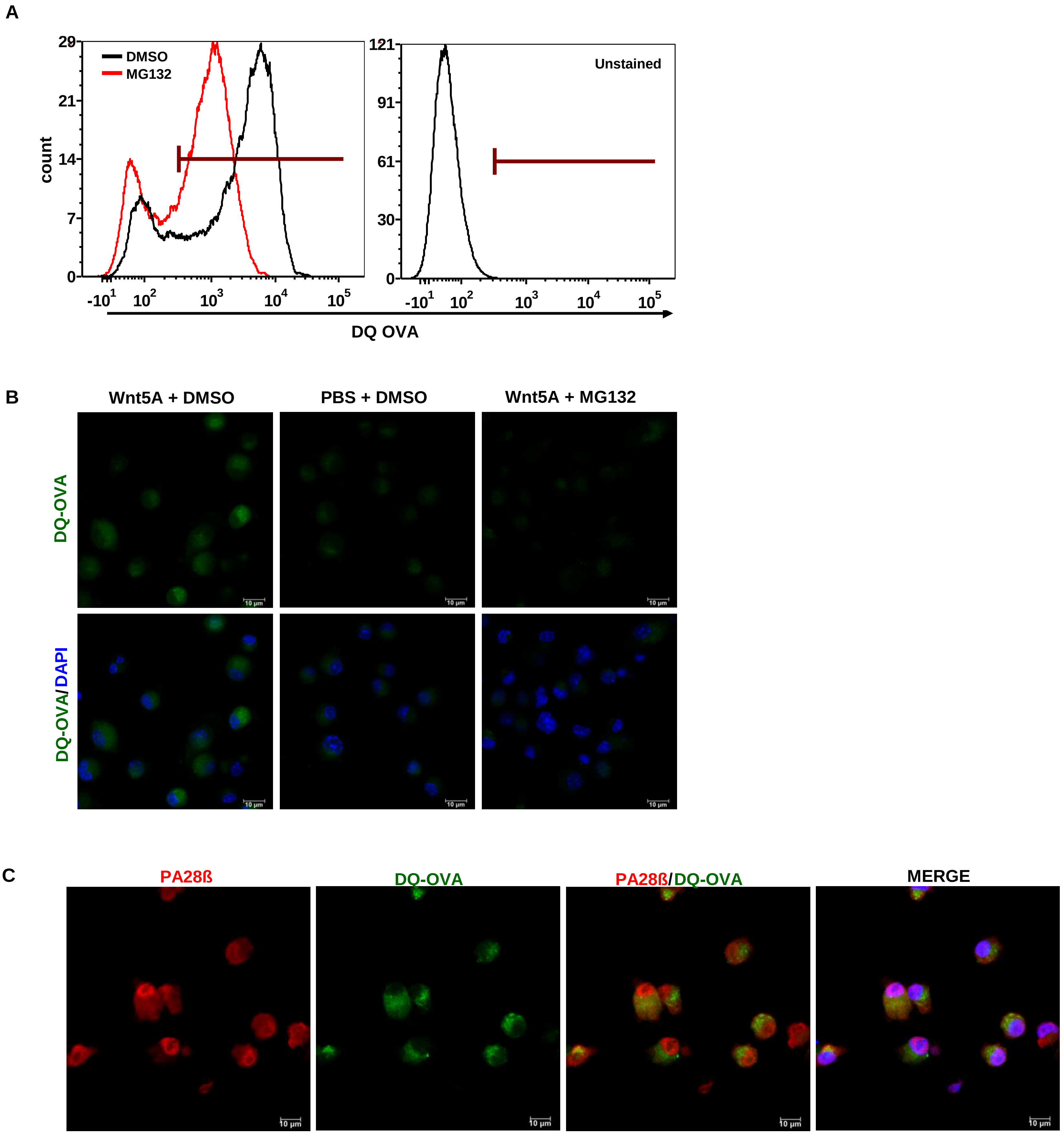


**Figure S4:** **<u>DQ-OVA processing in BMDC is proteasome-mediated:</u>** **A:** FACS demonstrating reduction in DQ-OVA fluorescence in BMDC after MG132 (proteasome inhibitor) treatment (10uM) for 30min as compared to control (DMSO: vehicle control treatment). **B:** Reduction in DQ-OVA intensity in presence of MG132 in Wnt5A treated BMDC as shown by confocal microscopy. **C:** DQ-OVA co-localizes with PA28ß (proteasome regulator) in DQ-OVA treated BMDC as demonstrated by confocal microscopy.

Supplementary fig: 5

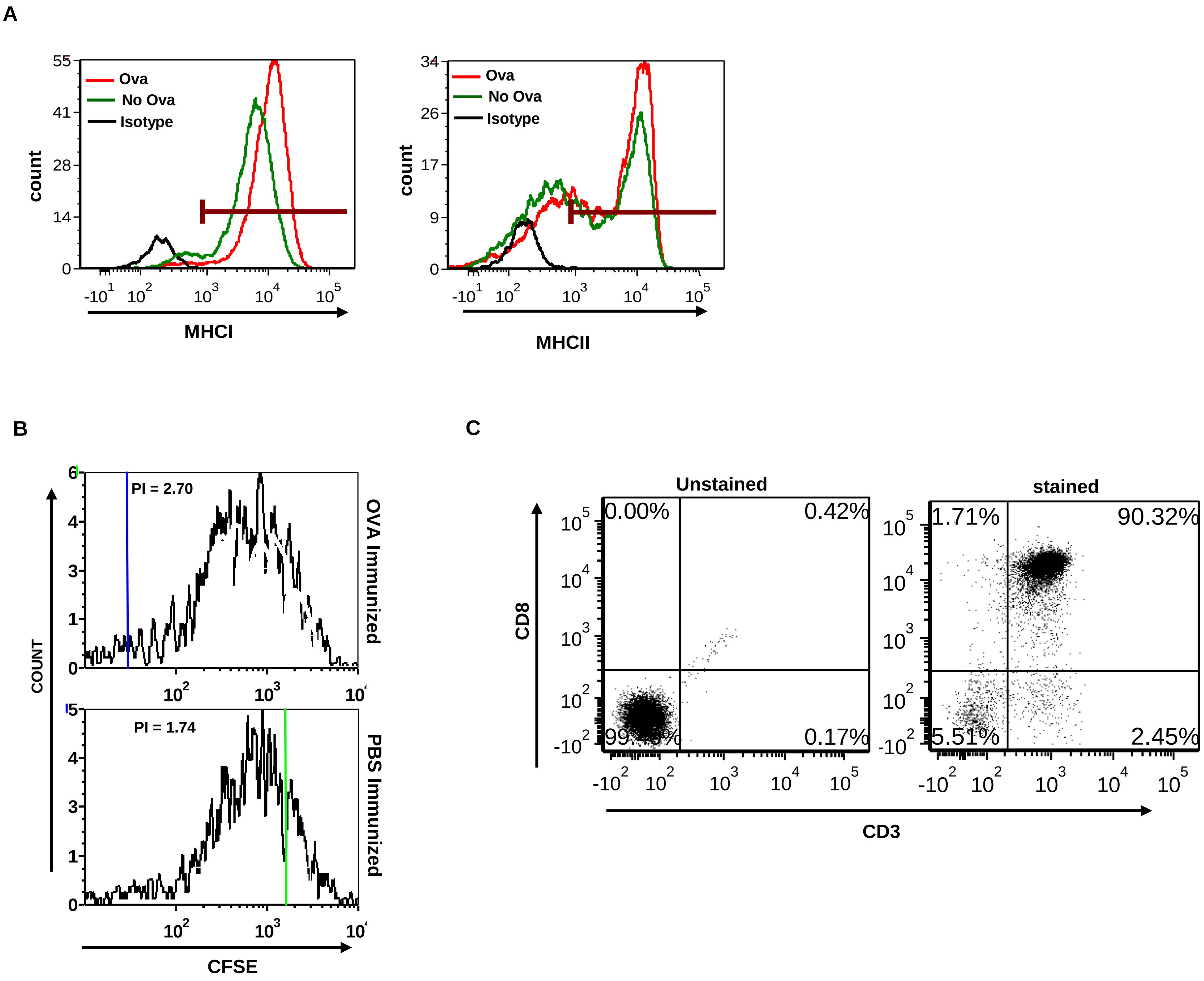


**Figure S5: <u>Immunization with OVA leads to strong recall response of OVA-specific CD8 T cells to OVA:</u>** **A:** FACS histogram showing increase in surface expression of MHC Class I and MHC Class II on BMDC after stimulation with 100ug/ml of OVA for 20 hrs. **B:** Representative FACS histogram demonstrating higher proliferative index (PI) of CD8 T cells from OVA immunized mice in response to OVA stimulation as compared to those harvested from mice where OVA was replaced by PBS, the vehicle control. PI calculation is explained in Materials & Methods. **C:** Representative FACS dot plot demonstrates CD8 T cell purity.
